# Observing changes in human functioning during induced sleep deficiency and recovery periods

**DOI:** 10.1101/690313

**Authors:** Jeremi K. Ochab, Jerzy Szwed, Katarzyna Oleś, Anna Bereś, Dante R. Chialvo, Aleksandra Domagalik, Magdalena Fafrowicz, Ewa Gudowska-Nowak, Tadeusz Marek, Maciej A. Nowak, Halszka Ogińska

**Affiliations:** Institute of Theoretical Physics, Jagiellonian University, Kraków, 30-348, Poland; M. Kac Complex Systems Research Center, Jagiellonian University, Kraków, 30-348, Poland; Department of Cognitive Neuroscience and Neuroergonomics, Jagiellonian University, Kraków, 30-348, Poland; Małopolska Center of Biotechnology, Jagiellonian University, Kraków, 30-348, Poland; Center for Complex Systems & Brain Sciences (CEMSC3), Universidad Nacional de San Martín, Buenos Aires, 1650. San Martín, Argentina; Consejo Nacional de Investigaciones Científicas y Tecnológicas (CONICET), Buenos Aires, C1425FQB, Argentina

## Abstract

The duration of sleep, wakefulness and dynamic changes in human performance are determined by neural and genetic mechanisms. Sleep deprivation and chronic restriction of sleep cause perturbations of circadian rhythmicity and degradation of waking alertness as reflected in attention, cognitive efficiency and memory. In this work we report on multiple neurobehavioral correlates of sleep loss in healthy adults in an unprecedented study comprising 21 consecutive days divided into periods of 4 days of regular life (a baseline), 10 days of chronic partial sleep restriction and 7 days of recovery. Throughout the whole experiment we continuously measured the spontaneous locomotor activity by means of actigraphy with 1-minute resolution in two acquisition modes (frequency and intensity of movement). Moreover, on daily basis the subjects were undergoing EEG measurements (64-electrodes with 500 Hz sampling frequency): resting state with eyes open and closed (RS; 8 minutes long each) followed by Stroop task (ST; 22 minutes). Altogether we analyzed actigraphy (distributions of rest and activity durations), behavioral measures (accuracy and reaction times from Stroop task) and EEG (amplitudes, latencies and scalp maps of event-related potentials from Stroop task and power spectra from resting states). The actigraphy measures clearly indicate rapid changes after sleep restriction onset, confirming our former investigations — the novel insight is a slow and incomplete relaxation to the original locomotor behavior. The pattern of partial recovery appears also in accuracy (in ST) and power of delta rhythm (in RS). The impact of sleep loss is also evident in reaction times (in ST), yet followed by complete recovery, and finally in ERP amplitudes and latencies, which however did not return to the baseline at all. The results indicate that short periods (a few days) of recovery sleep subsequent to prolonged periods of sleep restriction are overall insufficient to recover fully.

## Introduction

Optimal duration of sleep has been documented as fundamentally linked to health^1^ and cognitive performance^2^. Increasing evidence shows that sleep loss deteriorates basic neurobehavioral functions as working memory, attention or decision making, and contribute to negative health consequences related to circadian desynchrony^3^. Both partial (defined as a reduction in a sleep time over a 24-hour period, relative to individual sleep routine; also referred to as ‘sleep restriction’) and total (defined as a complete lack of sleep in a 24-hour period; also referred to as ‘acute’) sleep deprivation are linked with deficits in a cognitive performance^4^, higher risks of motor accidents^5,6^ and medical errors^7^. Furthermore, insufficient sleep is also associated with ill health, such as higher risk of diabetes, obesity, heart problems, and even stroke^8^. Partial sleep reduction to 6 (or less) hours per night for 14 consecutive nights results in comparable cognitive deficits to those observed after 1-2 nights of total sleep deprivation^4^, and include problems with attention and working memory, as well as high subjective levels of sleepiness.

Prediction of performance deficit and identification of biomarkers prognosticating dropping alertness connected to acute or chronic sleep loss can be a difficult task, because the magnitude of fatigue, sleepiness, and degradation of cognitive performance in response to acute and chronic sleep restrictions involves variability due to individual traits^9–11^. These can be related to chronotypes (“morningness” versus “eveningness”, i.e., a tendency to stay optimally attentive during a given time of day), which exhibit differences in sleep regulation and response to sleep fragmentation^12,13^, and which in turn may mask response to sleep deficit in individuals.

While problems linked with partial and total sleep deprivation are, however, fairly well reported in the literature, only a handful of studies looked at the processes involved in the recovery following an extended period of sleep restriction, and a majority of those that did, only looked at a very limited period of recovery^4,14–18^, typically between 1-3 days.

Therefore in this paper, we report the results from a longitudinal study that looked at the behavioral and neural effects of sleep restriction, as well as the recovery phase that followed. The study lasted for 21 days in natural enviroment, with the first 4 days of a baseline typical routine (BASE), followed by a 10-day period of partial sleep restriction (SR), and a week of recovery (RCV). Each day, participants took part in an electroencephalography (EEG) session, during which they did a classic Stroop task, and filled out a number of questionnaires aimed to measure their subjective levels of sleepiness and affect. The Stroop test is widely used to measure the cognitive functions such as attention, processing speed, cognitive flexibility^19^, and working memory^20^. It assesses the ability to inhibit cognitive interference, which occurs when the processing of a stimulus feature affects the simultaneous processing of another attribute of the same stimulus^21^.

As a result, it requires a considerable degree of concentration to execute it both fast and correctly, and therefore in numerous studies it has been used to test the effects of sleep deprivation on performance^22–25^.

Additionally, during the whole period of 21 days, participants’ locomotor activity was monitored with the use of actigraphy. The results primarily showing changes during sleep restriction in multiple measures: behavioral, actigraphic, and neuronal, are unanimous. Secondly, and even more importantly, their return to baseline during the long recovery period is not consistent – behavioral performance tends to revert to its normal level, whereas actigraphy measures do so only partly, and EEG activity shows no reversion.

The next section, Materials and Methods, describes the selection of subjects, design of the study, and data acquisition procedures, and detailed methodology for data analysis (including actigraphy, behavioral measures, and both task and resting state EEG). The Results section first reports on actigraphy, then it presents measures of performance in behavioral task, it continues with the changes in amplitude and latency of event-related potentials (ERP), and finally it provides the results of EEG power spectrum analysis. The results are summarised in Table 1 and followed by a thorough Discussion section. We further provide some Supplementary Figures and Tables, specifically for extended treatment of actigraphy, as well as detailed reporting on statistics of behavioral and EEG measures.

**Table 1.**
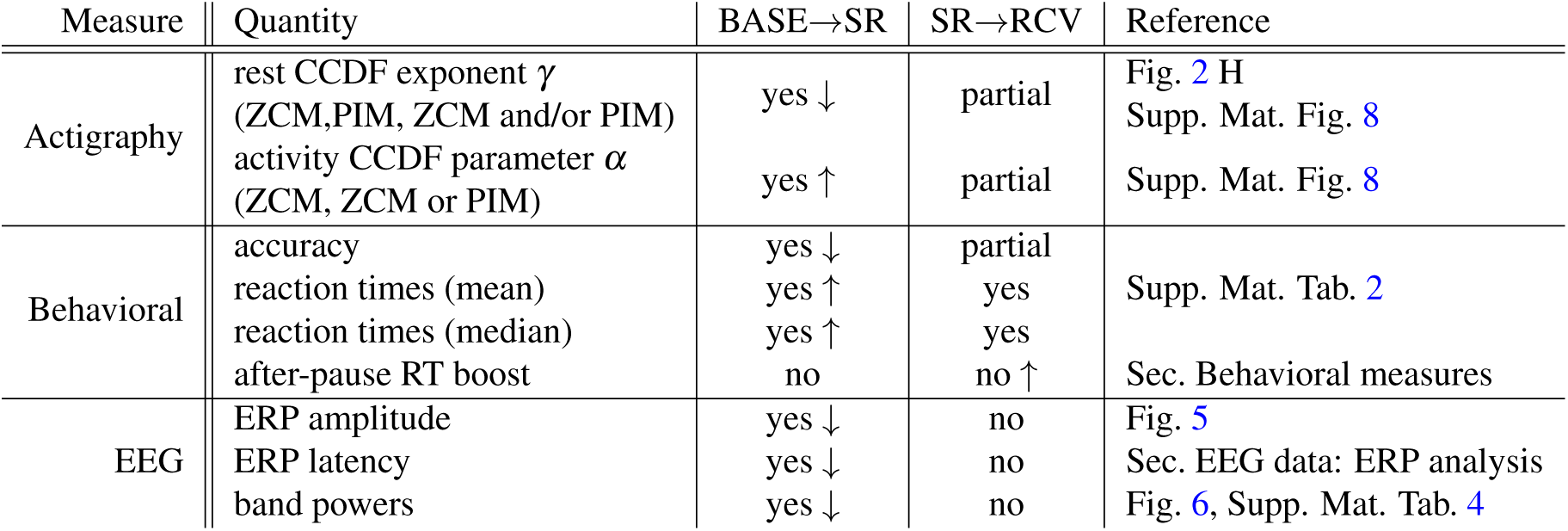
A simplified summary of the results where the difference between baseline and sleep restriction or recovery was found significant. The arrows *↑* and *↓* indicate increase or decrease of a given measure. Partial return indicates a tendency for a given measure to go back to baseline value, even if they still remain different.

## Materials and Methods

### Participants

The total number of 23 participants underwent our experiment. However, based on actigraphy recordings, 4 participants were removed from analysis due to failure to comply with the prescribed sleep restriction. The analyses of actigraphy take into account all the remaining 19 subjects (9 morning, 10 evening). Due to a revision of the Stroop task design, another 6 subjects are left out, with 13 subjects remaining in the reported EEG and behavioral analyses.

Further, individual days for a given subject were excluded as follows:

- if towards the end of baseline period a given subject slept as little as during sleep restriction period,
- if a given subject failed to comply with the sleep restriction towards the end of its period, we removed its further days from the analysis, but left the recovery data,
- if in the middle of recovery a subject slept as little as during sleep restriction, we removed individual days from recovery data.

The subjects were recruited based on Pittsburgh Sleep Quality Index^26^ and Epworth Sleepiness Scale^27–30^ questionnaires. All participants were healthy, drug-free (including alcohol and nicotine), and reported regular sleeping patterns with no sleep-related problems. The mean age of the 13 subjects was 21.5 ± 1.3 y.o.; they were 12 women and 1 man; they had 6 evening and 7 morning chronotypes (measured by morning-evening orientation and subjective amplitude, described with Chronotype Questionnaire^31^).

All of the subjects provided their written informed consent and were financially reimbursed for their time. The study was approved by the Bioethics Committee of the Jagiellonian University, Kraków, Poland.

### Data acquisition

#### Procedure

As illustrated in Fig. 1, the whole study lasted 21 consecutive days, which were divided into 3 sleep conditions: a 4-day period of unrestricted sleep (‘baseline’; hereafter abbreviated to BASE), then a 10-day period of daily 30% sleep reduction relative to individual sleep need (‘sleep restriction’; SR) followed by a 7-day period of unrestricted sleep again (‘recovery’; RCV). The subjects were instructed to conduct their normal daily routines during BASE; during SR they were individually prescribed a reduced sleep duration to follow; during RCV they were instructed to sleep without any restrictions. The subjects were additionally instructed to refrain from partying for the whole duration of the study and from caffeine consumption before visiting the EEG laboratory. The subjects were informed that their sleeping patterns will be monitored with actigraphy.

**Figure 1.**
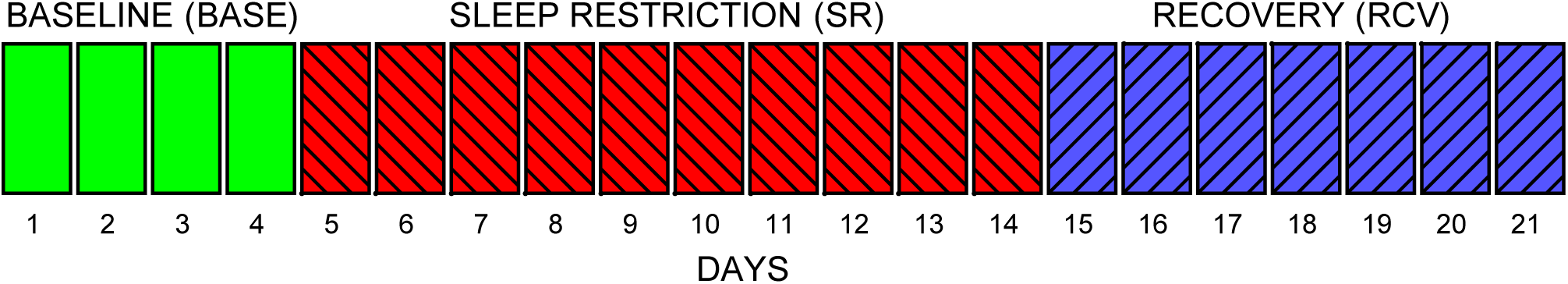
Study experimental design, showing number of days in each of the experimental sleep conditions.

Each day, participants’ brain activity was being monitored with an EEG (64 electrodes, Geodesic Sensor Net, EGI System 300, USA), while they were in a resting state (for 8 minutes with open eyes, RSeO, followed by 8 minutes with closed eyes, RSeC), and next performing Stroop test. The EEG session for a given participant took place in the laboratory at the same time every day, in the morning or evening according to the participant’s chronotype. Daily, before and after each experimental session, the severity of subjective sleepiness level was assessed by Karolinska Sleepiness Scale^32^ and participant’s mood was assessed with the use of Positive and Negative Affect Schedule^33^ questionnaire.

Additionally, the motor activity of participants was continuously recorded with actigraphy (Micro Motionlogger Sleep Watch, Ambulatory Monitoring, Inc., Ardsley, NY) in order to control their sleep timing and duration; a given participant wore the same device during the whole 21-day period. The actigraphs collected data in 1-min intervals in two modes: the zero-crossing mode (ZCM; i.e., frequency of activity signal crossing a zero threshold) and the proportional integrating measure (PIM; i.e., intensity or under the signal curve), see Fig. 2 A-E. The EEG recordings were performed in the laboratory, whereas the actigraphy data were collected continuously for the 21 days of experiment: both in the laboratory and in the natural environment.

**Figure 2.**
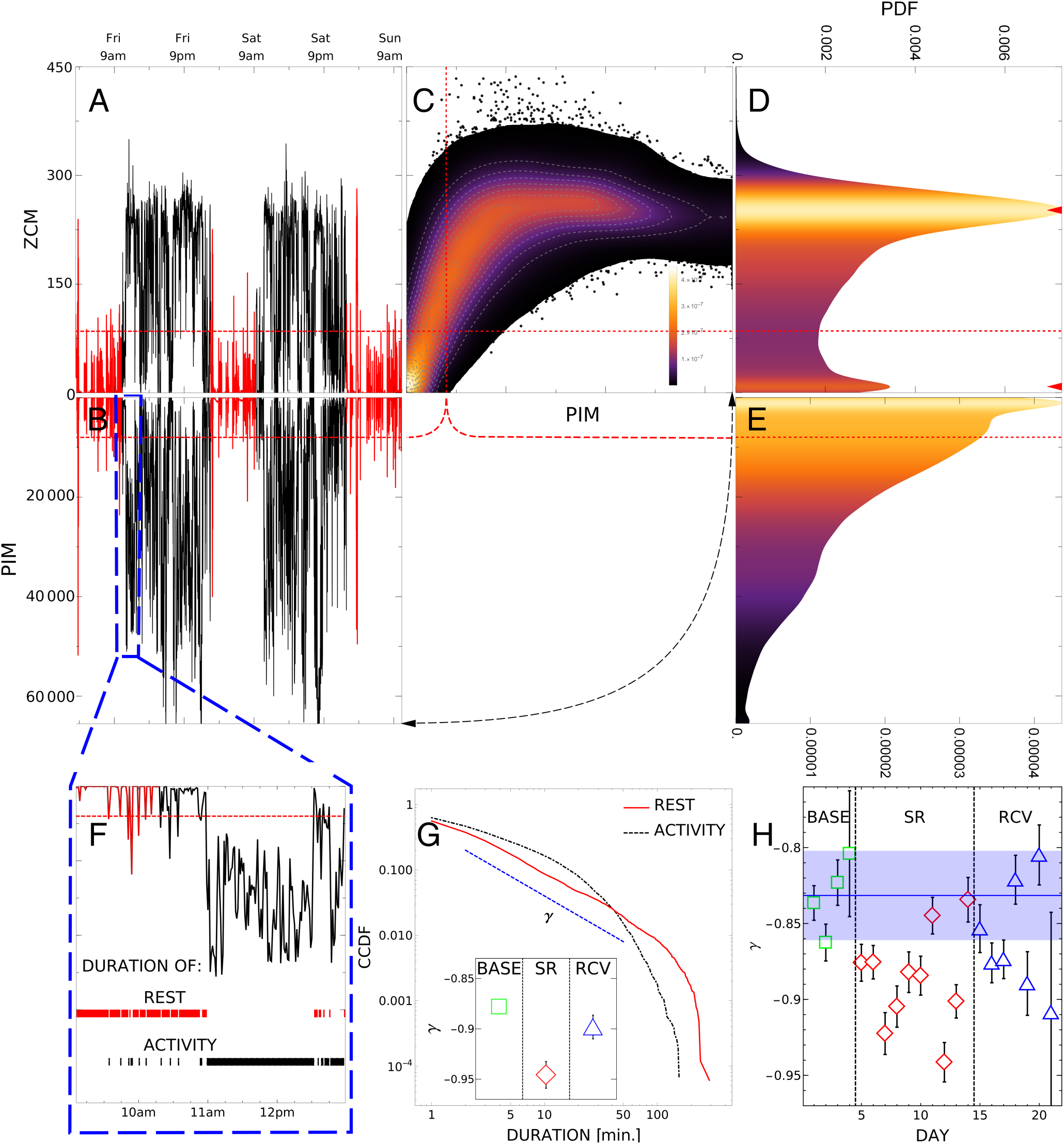
Summary of actigraphy analysis. A-B: two-day activity recording of a single subject in ZCM and PIM modes. C: density histogram of ZCM versus PIM activity (vertical and horizontal axes, respectively). D: histogram of ZCM recordings, with two peaks corresponding to diurnal and nocturnal activity marked by red triangles. E: histogram of PIM recordings. F: a sample of rest and activity periods defined by a threshold (red dashed line). G: complementary cumulative distributions of duration of rest and activity in log-log scale, with *γ* being the exponent of power-law tail of rest CCDF. Inset: *γ* exponents in baseline, sleep restriction and recovery conditions. H: *γ* exponents for each day of the experiment obtained from rest duration CCDF of all subjects.

#### Stroop task

Each day the subjects’ performance in terms of cognitive information processing was measured in a classic Stroop test. The subjects were asked to decide whether the name of the color matches with the ink in which it is written (congruent conditions) or not (incongruent conditions). So the more automated task (reading the word) interfered with the less automated task (naming the ink color) resulting the difficulty in inhibiting more automated process known as the Stroop effect. In the final version, the test was designed with four different colors (red, blue, green, and yellow) and only that data is reported here. During the first stage of the experiment (6 participants, not reported here) the test design included more colors but was revised to involve less variability. In total there were *N*_*s*_ = 432 randomly shuffled stimuli presented in 3 separate blocks (144 stimuli each) with a short break in between each block, as indicated in Fig. 3. Half of the stimuli were congruent, and half were incongruent. The inter-stimulus interval between each stimulus was between 1500 ms and 3500 ms (in steps of 400 ms, with 2500 ms on average); the entire task lasted 22 minutes and 11 seconds on average. Behavioral measures of all stimuli were analyzed, even if the corresponding epochs in EEG data were removed due to insufficient quality.

**Figure 3.**
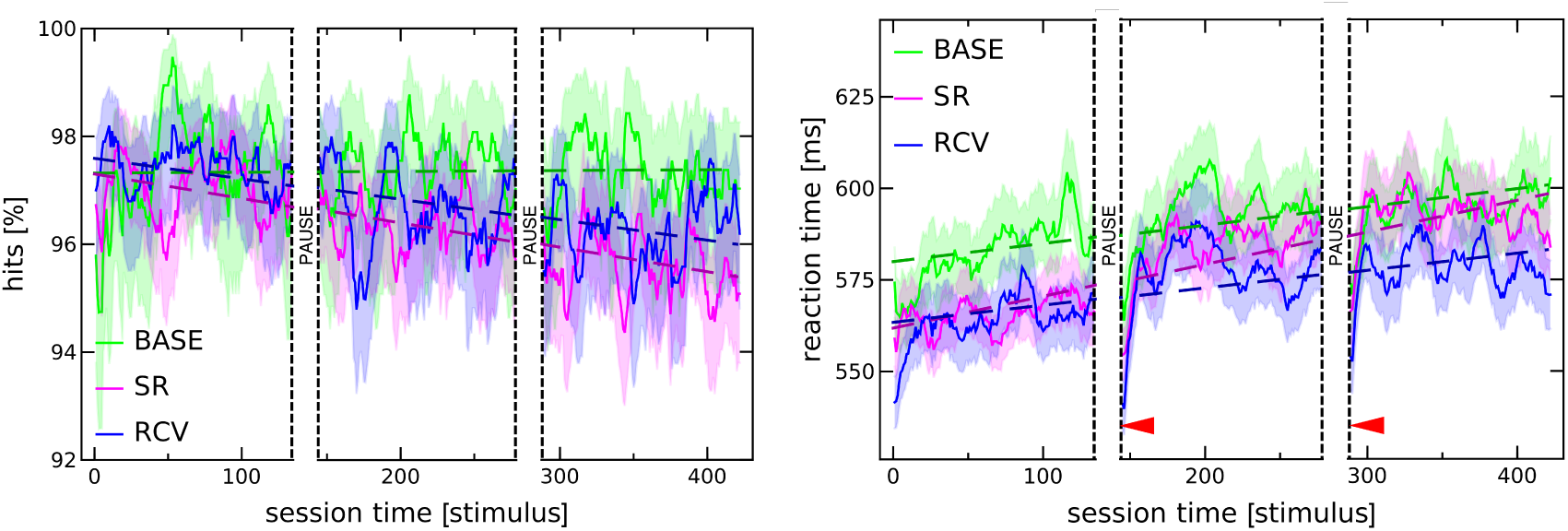
Changes in accuracy (left) and reaction times (right) exhibit trends as functions of time spent on performing the task (measured by the number of stimuli presented). The continuous lines represent means over subjects and days within a given condition, and the shaded regions 95% confidence intervals; dashed lines are linear fits. Vertical dashed lines indicate pauses between blocks of the Stroop task. Detailed description is in the text below.

All participants underwent one training session before beginning of the experiment in order to avoid the learning effect.

### Data analysis

Whenever possible, when reporting on statistical significance of a result we provide (in the text or in the Supplementary Material) a bare p-value. Unless otherwise explicitly stated we use the term “significant/insignificant” for significance level *α* = 0.05 with multiple-comparisons taken into account (Holm-Bonferroni method with three comparisons: BASE-SR, BASE-RCV, and SR-RCV). Particular methods and tests depend on data type, and are described in detail below.

#### Actigraphy

We followed the procedures presented in the detailed actigraphy analysis of sleep deprived individuals in^34,35^ (on a different data set). First, the raw actigraph data *X* (*t*), see Fig. 2 panels A and B, were segmented into 18-hour days (4 hours sleep followed by 14 hours wakefulness); data which did not fit into these durations or contained other artifacts (5.2% of all data) was discarded. The fixed length removes statistical artifacts connected with inter- and intraindividual variance of sleep duration and the forced difference between restricted and unrestricted sleep. Next, based on a predetermined threshold, the data was split into ‘activity’ and ‘rest’ (data points above and below the threshold, respectively), see Fig. 2 panel F. Lastly, their durations (lengths of uninterrupted periods of activity or rest) were extracted. The thresholds were set to *T*_*ZCM*_ = 85 and *T*_*PIM*_ = 8000, based on the optimal goodness of fit of a power law (3) to the distribution of rest durations.

Since the ZCM and PIM actigraphy modes may contain complementary information (basically, the frequency and intensity of movements), in the present paper we added a new methodological element analyzing four scenarios:

- threshold only for ZCM, see panel A in Fig. 2,
- threshold only for PIM, see panel B,
- jointly: when *both* ZCM *and* PIM cross a threshold, see panel C,
- jointly: when *either* ZCM *or* PIM crosses a threshold, see panel C.

For a given scenario, we counted the number of activity/rest periods of given duration time jointly for all subjects, and calculated the resulting probability density function (PDF) *p*(*τ*) of duration time *τ*. To better assess statistics of rare events in tails of PDF we construct, as the main measure of discussed phenomena, the complementary cumulative distributions (CCDF) *C*(*a*) of durations *a*, see Fig. 2 G:

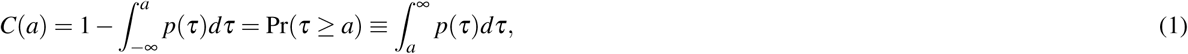

which represents surviving probability for the system to stay in a given state for up to the time *a*. The function *C*(*a*) is equal to one for a minimal value *a* = 0 and tends to zero for *a →* ∞. For a stationary time series the survival probability *C*(*a*) is expected to have a characteristic scale (relaxation time *τ*_*rel*_) related to the probability per unit of time *λ* to undergo the change of the state. The rate *λ*, named the hazard rate in the theory of critical phenomena, denotes then the ratio

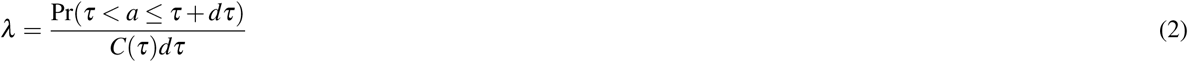

and can be associated with *C*(*a*) representing a simple exponential function of dwell times, *C*(*a*) = *e*^*−λa*^ with *λ* = 1*/τ*_*rel*_. As discussed elsewhere^34–36^, in case of disordered systems, the notion of the survival function and the corresponding (hazard) rate *λ* (*τ*) can present much more complex behavior, strongly deviating from a simple Poissonian-like occurrence of the ‘threshold-exceeding’ events.

For the purpose of statistical analysis, the numerical estimates of cumulative distributions was fitted with two mathematical formulae, power-law of the form

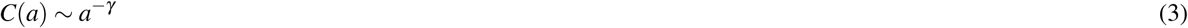

for rest periods and a stretched exponential

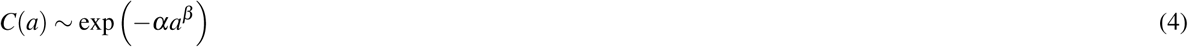

for activity periods.

For robustness, the cumulative distributions were bootstrapped (on the BASE/SR/RCV period level there were 1000 samples with 60 daily recordings each; on the day-by-day level there were all combinations of *N*_*d*_ *−* 2 daily recordings, ranging 10 *−* 171 samples a day, where *N*_*d*_ is the number of available recordings for a given day). The ordinary least-squares fitting was performed on log-log or log-linear data, respectively for a power-law and stretched exponential, in order to account for the tails in the distributions. The fitting was weighted, with weights coming from bootstrap standard deviations, which removed variance caused by the very end of the distribution tails. The fitted parameters *α, β* and *γ* were then compared for several period and subject combinations. The error bars in Fig. 2 G-H are 95% confidence intervals of the respective fit. The statistical significance of differences between exponents *γ* was tested by t-test^37,38^ for differences in linear regression slopes (version with unequal residual variances; the test is based on fit residuals, which in this case are very highly non-normal due to the very end of the distribution tail, cf. rest distribution in Fig. 2 G; to fulfil normality assumption we took approximately only the first half of residuals).

#### Behavioral measures

The two measures scrutinized in this paper are: accuracy (percentage of correct responses) and reaction times, for results see Sec. Stroop test – time-on-task effect. The curves shown in Fig. 3 are averages over subjects and days within a given condition (baseline, sleep restriction, recovery) computed over overlapping moving windows of *w* = 10 consecutive stimuli.

Since the responses have binary values (correct or error) and the percentage of hits is high (around 96%), the accuracy has a highly skewed binomial distribution, and so we use Clopper–Pearson binomial proportion 95% confidence intervals. For reaction times, we use standard 95% CI.

In the figure, the dashed lines represent fitted linear models. The linear regression did not take into account the pauses in the experiment (nor the first 6 stimuli in each block for RTs) i.e., the input data were only the 432 (414) stimuli responses, however, in this case we did not use window averages over consecutive stimuli. The change in accuracy and reaction times reported in Supp. Mat. Table 2 is simply the slope (together with its 95% CI computed from the standard error of the fit) of the regression line multiplied by *N*_*s*_ *−* 1. The statistical significance of differences between the slopes can be assessed simply by investigating the confidence intervals, but we also provide p-values from the appropriate t-test^37,38^ (version with unequal residual variances). One must be aware, however, that in the case of accuracy and mean reaction time the residuals do not fulfill normality assumption. Since reaction times have a skewed distribution, see Fig. 9 in Supp. Mat., in Table 2 we also report on their medians, in which case the fit residuals are normal.

**Table 2.** Average changes in accuracy and reaction times between the beginning and end of an experimental session (432 stimuli, 22 minutes) computed from linear regression slopes. Beside means (or medians) and 95% CI for each condition on the left, we also report on p-values for pairwise differences between conditions (e.g., the element in the 1st row and 2nd column gives the p-value for BASE-SR). These results are summarized in Sec. Behavioral measures.

| ACCURACY |  |  |  |  |  |
| --- | --- | --- | --- | --- | --- |
|  | mean [%] | 95% CI | BASE | p-values<br>SR | RCV |
| BASE | -0.062 | [-0.76, 0.63] | – | $6.5 \times 10^{-6}$ | **0.00019** |
| SR | -1.96 | [-2.53, -1.38] |  | – | 0.23 |
| RCV | -1.64 | [-2.26, -1.01] |  |  | – |

| REACTION TIMES (means) |  |  |  |  |  |
| --- | --- | --- | --- | --- | --- |
|  | mean [ms] | 95% CI | BASE | p-values<br>SR | RCV |
| BASE | 21.3 | [16.2, 26.5] | – | $1.8 \times 10^{-6}$ | 0.37 |
| SR | 37.5 | [32.8, 42.1] | | – | $1.3 \times 10^{-7}$ |
| RCV | 20.2 | [15.4, 25.0] |  |  | – |

**Table 2.** Average changes in accuracy and reaction times between the beginning and end of an experimental session (432 stimuli, 22 minutes) computed from linear regression slopes.
| REACTION TIMES (medians) |  |  |  |  |  |
| --- | --- | --- | --- | --- | --- |
|  | median [ms] | 95% CI | BASE | p-values<br>SR | RCV |
| BASE | 17.3 | [12.0, 22.6] | – | 0.028 | 0.44 |
| SR | 23.4 | [19.7, 27.1] |  | – | 0.020 |
| RCV | 17.8 | [13.8, 21.9] |  |  | – |

To measure the after-pause decrease in reaction times, the linear fit described above was subtracted from them. To test the differences in depth of the decrease between conditions all individual responses to the first 6 stimuli in the 2nd and 3rd block were used. Since their distribution is non-normal and the variances are not equal Mann-Whitney test was performed.

#### EEG

The EEG analyses were performed with the use of EEGlab v. 13^39^ and ERPLAB 6.0^40^ on MATLAB R2016b. The time series were digitally filtered (0.5–35 Hz), the reference was recomputed from Cz electrode to average, bad channels and time epochs with artefacts were removed. An Independent Component Analysis (ICA) was used to separate and remove physiological artefacts, including saccade-related spike potentials^39^. We report in detail on electrodes *vref*, 4, and 34 from EGI’s 64-channel system, which correspond to Cz, Pz, and FCz in 10-10 system within 1 cm accuracy.

The power spectra of continuous EEG signals in resting state were computed for chosen electrodes using Welch’s overlapped segment averaging estimator after the above pre-processing. The same procedure was repeated for epoched EEG recordings when subjects performed Stroop task. Powers were then computed separately for each subject on each day in four sub-bands: delta (1-4 Hz), theta (4-8 Hz), alpha (8-13 Hz), and beta (13-30 Hz). Their distributions in each band and sleep condition (BASE, SR, or RCV) were in general non-normal (the highest p-value 0.066 in Shapiro-Wilk test was obtained for beta band in baseline period; others were orders of magnitude smaller). Consequently, instead of ANOVA we used Kruskal-Wallis test^41^ followed by Conover-Iman post hoc^42^ (chosen instead of the usual Dunn’s test for its reportedly greater power) as implemented in R^43^ to test for stochastic dominance of the samples.

The event-related potentials were measured with segments from 200 ms before to 1000 ms after the onset of the target stimulus, with the pre-stimulus baseline subtracted. The potentials are grand averages for a single condition (BASE/SR/RCV) and stimuli type (congruent/incongruent). In total there were 32 recordings for baseline, 47 for recovery and 73 for sleep restriction, but in SR period we show only 38 sets from days 10 to 14 – the reason being on one hand a more uniform sample size, and on the other stronger expected effects in the later phase of sleep deficit. Since, as reported in in^44^, there was no significant interaction between the sleep condition and stimuli type, subsequently the ERPs from congruent and incongruent stimuli were averaged for robustness and a more concise presentation in Fig. 4-5. The significance tests presented in Fig. 5 were performed at each time point separately with a t-test or Mann-Whitney test, depending on which test assumptions were met.

**Figure 4.**
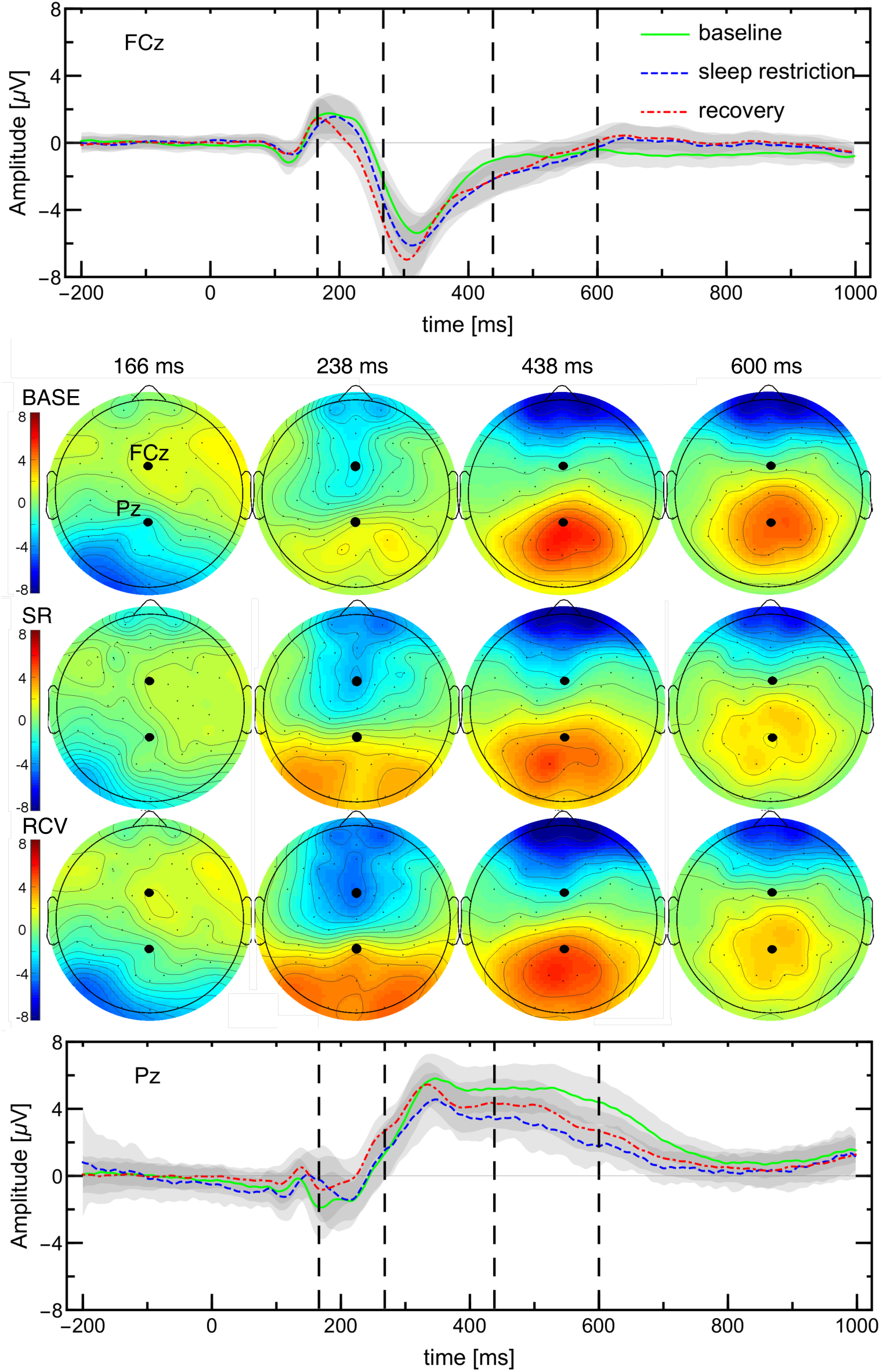
Activation maps and ERPs in response to Stroop task (average of congruent and incongruent stimuli). The maps are shown in columns corresponding to four chosen time points (also indicated by dashed lines) and ERPs are shown at the two locations (Pz and FCz) that correspond to the largest differences between baseline and SR (days 10-14) or RCV periods. The amplitude scale in the maps is the same as in the ERP plots.

**Figure 5.**
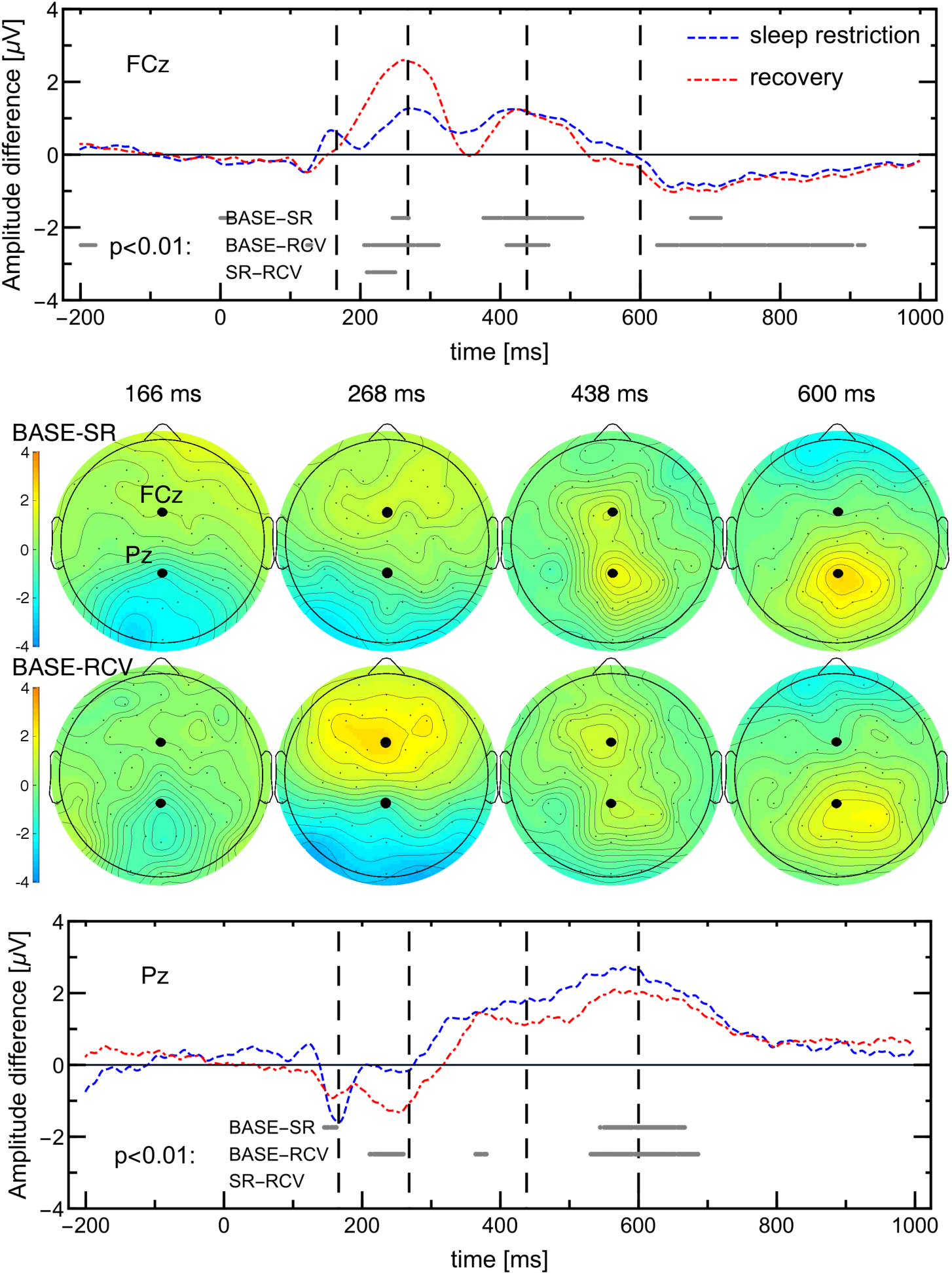
Differences in ERP amplitudes between BASE and SR or RCV periods, derived from Fig. 4. Note that the scale is twice as small as therein. Grey lines in the lower part indicate time points at which t-tests for pairwise comparisons yielded value *p <* 0.01.

**Figure 6.**
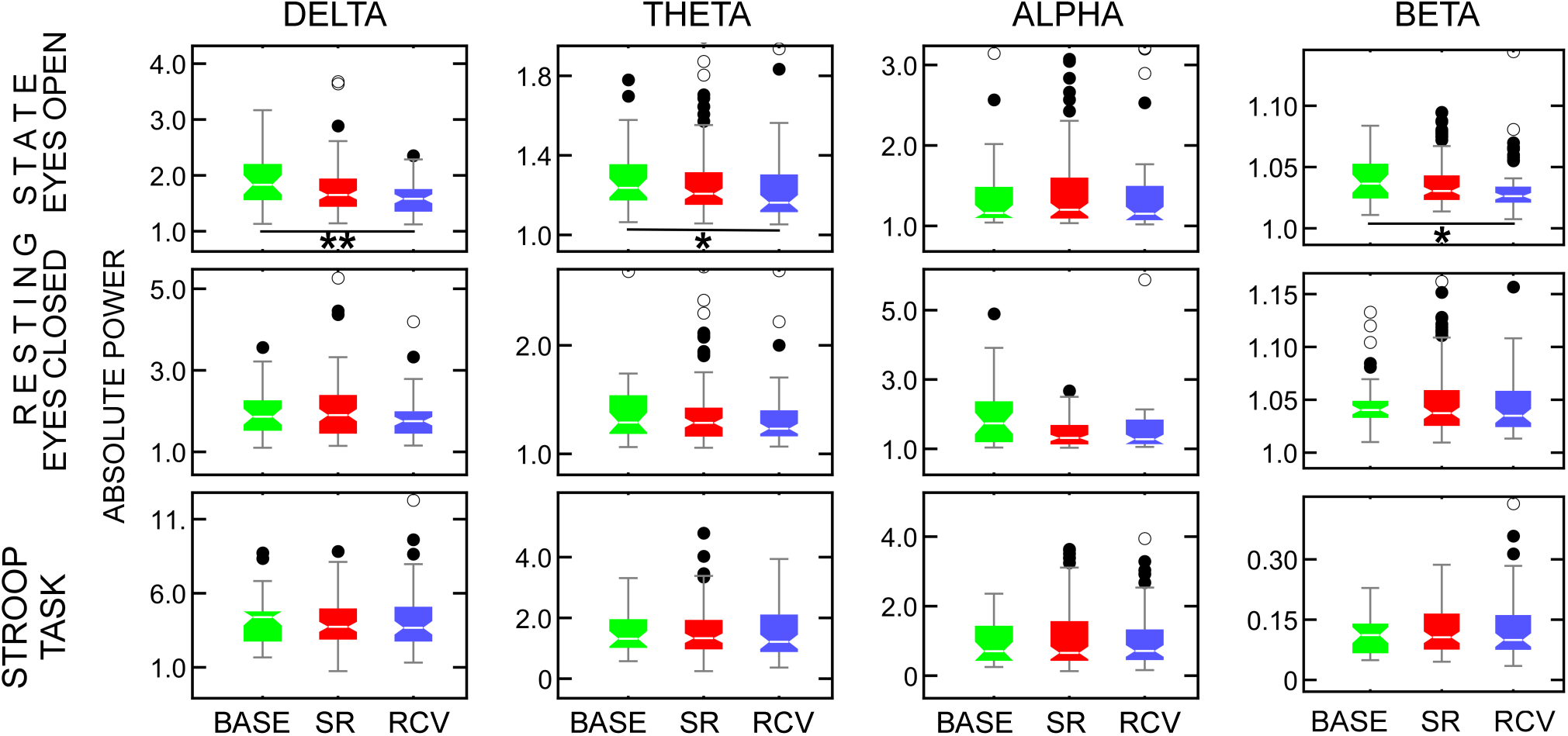
The absolute power of *δ* (1-4 Hz) (first column), *θ* (4-8 Hz) (second column), *α* (8-13 Hz) (third column) and *β* waves (13-30 Hz) (fourth column) for resting state with eyes open (top row), eyes closed (middle row), and for Stroop test (bottom row) computed at Cz electrode. The boxes show median and interquartile range, together with near (full circles) and far (empty circles) outliers. Some extreme outliers are not shown to retain the scale for visibility. Significant differences between conditions are marked with: * *p <* 0.05, ** *p <* 0.01.

**Figure 7.**
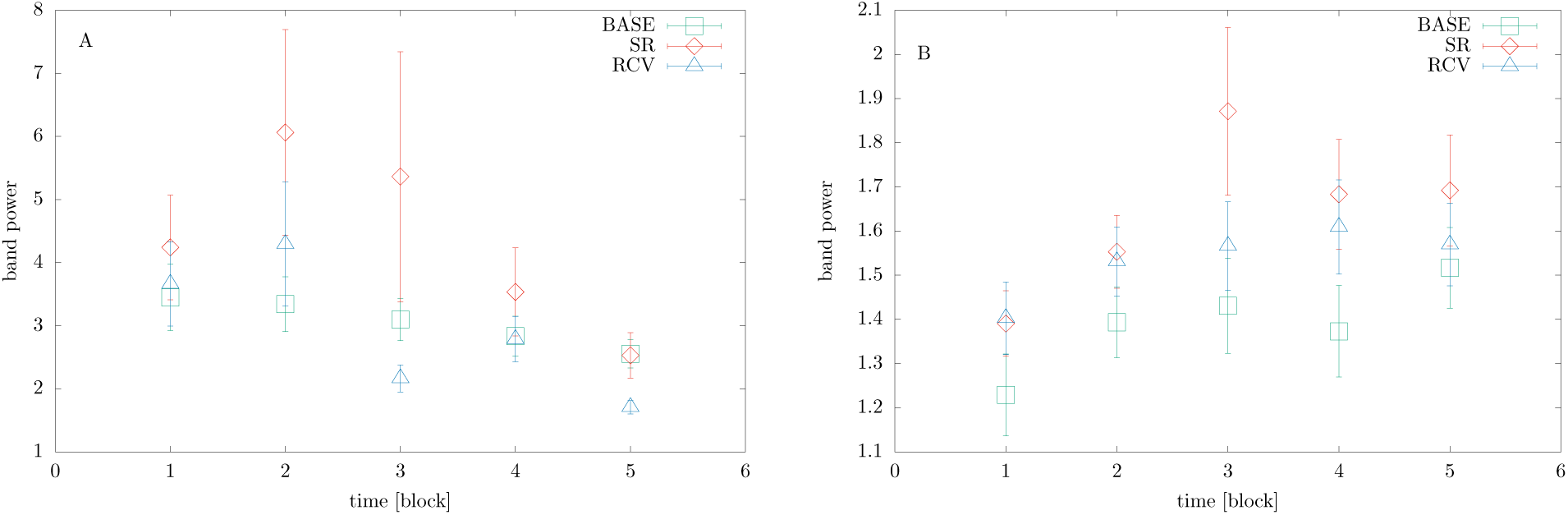
The power spectrum of alpha waves changing in time for resting state with eyes closed (left panel) and eyes open (right panel). Data from electrode Cz.

The peak time shifts between conditions was computed by minimizing difference between ERP curves on the [200ms, 300ms] interval.

#### Data availability

The datasets generated during and/or analyzed during the current study are available from the corresponding author on reasonable request.

## Results

### Actigraphy

Figure 2 summarizes actigraphy data characteristics and processing. A sample of raw ZCM and PIM time series is shown in panels A-B. As visible in panel C, the dependence between the two modes is linear only for small values. The resulting distributions of activity as recorded in ZCM and PIM (panels D-E, flipped to the side) also differ: ZCM has two clear peaks corresponding to daily activity and sleep, while PIM distribution monotonically falls down from low to high intensities. Panel F shows periods of activity and rest for PIM (respectively above and below *T*_*PIM*_ = 8000) obtained from an exemplary recording.

First, the activity period durations undergo stretched exponential distributions as found in our previous papers^34,35^. In this experiment we found some dependence of the exponent *α* on the measurement period, contrary to our previous results. This may be due to different experimental conditions (21 consecutive days as contrasted to two separate 7 days periods). The exponent *α* in 4 different mode combinations can be found in Fig. 8 in Supp. Mat.

**Figure 8.**
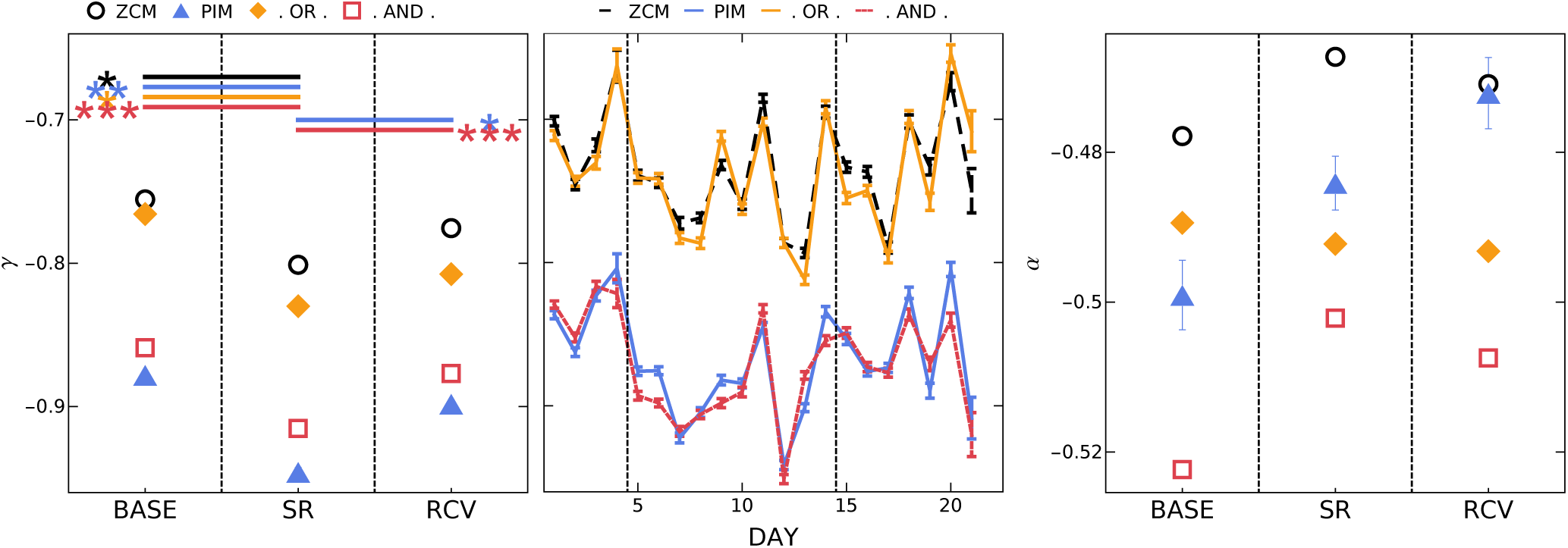
(Left panel) The gamma exponent of cumulative distribution *C*(*a*) of rest periods (defined as below ZCM, PIM, ZCM **or** PIM threshold, ZCM **and** PIM threshold) in the 3 measured conditions: baseline, sleep restriction and recovery, and (middle panel) as function of consecutive days. Asterisks denote significance of differences between conditions, color coded for the respective rest thresholds. (Right panel) The alpha exponent of cumulative distribution *C*(*a*) of activity state (defined as above respective thresholds)

More importantly, we concentrate on the CCDF of resting period durations *C*(*a*). In Fig. 2 G-H, we present the *γ* exponents (in the PIM mode at the group level) as a function of the measuring period and as a function of consecutive days. Consistently with our earlier work^34,35^ the distribution’s slope changes rapidly once the sleep loss appears (statistically significant with *p* = 0.0017).

The new result is long recovery time: even one week of regular sleep does not stabilize the distributions (the difference between SR/RCV is significant, *p* = 0.019, but smaller, while for BASE/RCV it is still visible but below confidence threshold, *p* = 0.16). Similar effect, although exhibiting more variability than in the PIM mode, is observed in the ZCM mode, cf. Fig. 8 in Supp. Mat.

Having at our disposal two different measuring modes we posed the question which mode or combination of modes is more informative of the changes resulting from sleep loss. The question follows from the observation that both modes are correlated but the relation is not linear, Fig. 2 C. Consequently, the division into resting and activity periods can differ in both modes. Thus, as indicated in Sec. Data Analysis: Actigraphy, we additionally tested exponents of distributions generated by the two modes simultaneously: either by defining resting states by alternative of ZCM and PIM thresholds (below ZCM or below PIM) or their conjunction (below ZCM and below PIM), see Fig. 8 in Supp. Mat. Comparison of the two measurement modes and their combinations shows that the use of both modes does not enhance the result and the PIM measurement mode seems to be most suitable for this particular experiment.

All the above combinations show a general trend – a significant change of the exponent during sleep restriction period and a partial return towards baseline values.

### Stroop test – time-on-task effect

The sleep restriction period influenced participants cognitive functioning as tested with the Stroop task. The main observation is poorer behavioral performance on the Stroop test during sleep restriction period followed by a gradual, incomplete recovery^44^. There was a tendency to have fewer correct responses during sleep restriction period -– mainly due to a significantly higher rate of errors and omissions, and a slightly increased rate of double responses. A decreasing tendency in reaction times (RT) for the first few days of the experiment was also observed, which can be interpreted as learning the Stroop task. As expected^45^, the distributions of RTs for congruent stimuli have smaller means and variances than for incongruent ones. Somewhat counter-intuitively, for incongruent stimuli there are more correct responses in the entire experiment than for congruent.

Here, we focus on how the time spent on performing Stroop task (within a given experimental session) affects accuracy and reaction times, see Fig. 3. The technical details on statistical computations are given in Sec. Behavioral measures.

The accuracy of responses, systematically and significantly decreases during session for SR and RCV conditions (by 1.96% and 1.64%, respectively), but not for the baseline period. The difference in the slopes between BASE and either SR or RCV is highly significant (*p <* 0.001; see Tab. 2 for confidence intervals and p-values). In other words, subjects undergoing and recovering from sleep restriction found it harder to maintain perfect accuracy during the 20-minute experimental session.

The change is more pronounced for reaction times. During a given session the RTs slightly increased, as visible in Fig. 3 (right). The difference between the last and first stimulus in a session, as given by linear regression, is 21.3 ms, 37.5 ms, 20.2 ms for BASE, SD and RCV conditions, respectively. As already visible, the difference between BASE and RCV is not significant, but it is for BASE/SR and for RCV/SR (*p <* 0.001; see Tab. 2 for details).

It should be noted, on one hand, that for accuracy and mean RTs the normality assumption on regression residuals is not fulfilled, and so the reported p-values for slope differences should be treated with caution. On the other hand, the confidence intervals obtained from the fitting do corroborate the conclusions. Additionally, due to high skewness of reaction time distributions^45^ (see Fig. 9 in Supp. Mat.), means might not be appropriate estimators, hence in Tab. 2 we also include results for median (where both differences for BASE/SR and RCV/SR are below *α* = 0.05 significance level, but marginally do not survive Holm-Bonferroni correction). We nevertheless believe that mean RTs – thanks to their sensitivity to skewness and outliers – can actually carry more information on atypical behaviour induced by sleep loss than medians.

**Figure 9.**
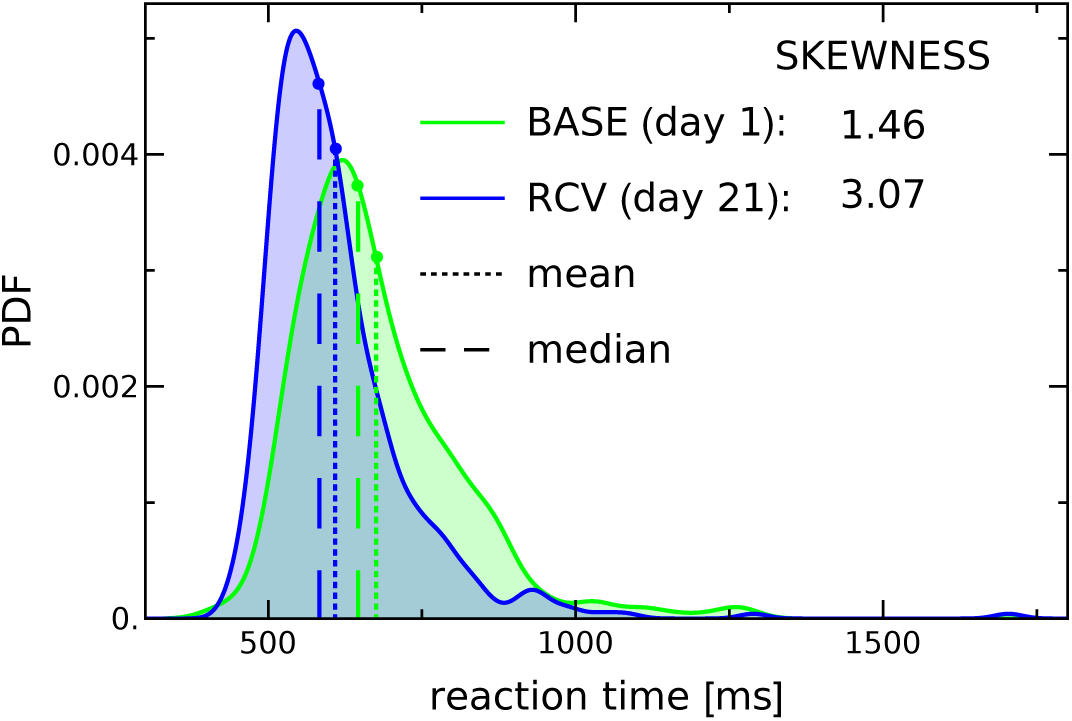
Histogram of reaction times for a single subject on the first and last day of the experiment. The distributions are highly skewed to the right, which results in significant differences between the mean and median.

In the Fig. 3, there are two dips in the RT curve: in one-third and two-thirds of the session, which is due to pauses between blocks of Stroop task (during which the electrodes were watered, and consequently the subjects were able to rest). These after-pause dips basically reset the reaction time to what it was at the beginning of the session. They take 6-16 consecutive stimuli, and are on average are 21.6, 22.2, and 34.3 ms deep for BASE, SR, and RCV, respectively. The difference between BASE and RCV is significant (*p* = 0.000016), but for SR and RCV it is not (*p* = 0.26, counter-intuitively large when compared with averages, which is due to high variance of individual responses).

In summary, the ratios of accurate responses and omissions directly follow the schedule of sleep restriction; they go back almost to normal already after the first day of recovery period. The difference between conditions in linear growth of reaction times (and linear decrease of accuracy) indicates that the sleep deprived subjects get tired more quickly. A difference between conditions in the after-pause drop in reaction times indicates that the pause gives more rest to recovering subjects. An interesting general observation is that the variability of response accuracy does not average out as well as the variability of reaction times.

### EEG data: ERP analysis – scalp maps and electrode dependence

The effects related to task performance in different conditions can be observed in event-related potentials.

It was observed that the P300 neural response was attenuated during and after sleep loss as compared to the baseline ^44^. Here, we analyses the whole scalp maps of ERP amplitudes to discovere how the brain response changes during different sleep condition.

The two individual electrodes shown in Fig. 4-5 were chosen by the maximal absolute difference between baseline activity and either SR or RCV period activity: for BASE-SR it was electrode Pz and for BASE-RCV it was electrode FCz. The times (166, 268, 438, 600 ms) at which the presented maps are plotted were also chosen as the times when the maximal differences occurred. The column of maps corresponding to the given time point is connected by dashed lines to that point in the ERP plot.

The general observation from Fig. 4 is that post-stimulus activations are mostly bipolar with strong positive activations in occipital and centro-parietal regions. Figure 5 additionally shows where and when the differences occur: visibly the baseline activations are greater at first in the frontal, then central, and finally parietal regions. Also visibly subjects in SR and RCV periods had stronger occipital activations earlier and the parietal activations disappeared sooner. Note that the greatest differences in ERP curves occur around and not at their peaks, which rather suggests a delay between them. The shift in time from the baseline ERP curve for Pz and FCz in Fig. 4 is: -2 ms and -12 ms, respectively, for SR and -20 ms and -26 ms for RCV.

### EEG data: power spectrum analysis

The absolute power of the delta, theta, alpha and beta waves were measured (Fig. 6) from EEG recordings during resting state (with eyes open and closed) and Stroop task. The power spectrum was computed at Cz electrode.

The highest power of the *δ* and *θ* oscillations were observed in the Stroop task. In contrast, the power of *β* waves during the task was drastically low. As expected, the *α* rhythms were significantly higher when subjects had their eyes closed when compared to performing the task or resting state with eyes open.

The shapes and variances of the band-power distributions differ between groups.The Kruskal-Wallis tests for differences between BASE, SR, and RCV periods yielded statistically significant results only for resting-state condition with eyes open, see Supp. Mat. Table 3 The differences were discovered between **BASE** and **RCV** in **delta, theta** and **beta bands**, see Fig. 6 upper panel and Supp. Mat. Table 4. On closer look, they came from a weak negative correlation with time, i.e., gradual decrease of power in these bands as the experiment progresses from BASE phase, to SR, and to RCV, which is visible in the top row in Fig. 6. The behavior of alpha band in resting state condition with eyes closed also did indicate differences between the periods, which were however below statistical significance.

**Table 3.**
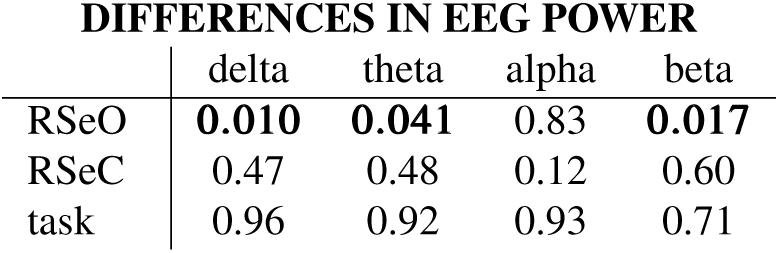
P-values of Kruskal-Wallis test with BASE, SR, RCV conditions as described in Methods: EEG section and Results: EEG data: power spectrum analysis.

**Table 4.** P-values of Conover-Iman post hoc test following Tab. 3.

|  | SR-RCV | BASE-SR | BASE-RCV |
| --- | --- | --- | --- |
| DELTA | 0.22 | 0.14 | **0.0068** |
| THETA | 0.13 | 0.62 | **0.046** |
| BETA | 0.12 | 0.36 | **0.015** |

We also examined how brain waves, in particular alpha waves, changed in time during the resting state part of the experiment. Figure 7 illustrates the alpha rhythms in both resting states (eyes open and closed) divided into 5 time blocks (96*s* each). This plot clearly presents two effects: firstly, the highest power for the sleep restriction period; and secondly, growing values of absolute power with the time of eyes being open, Fig. 7 B. These slow changes in brainwave power result in higher variance (visible in Fig. 6) and thus it is advisable to test for differences on time-dependent powers. What we found intriguing, is the values of alpha power in both sleep restriction (SR) and recovery periods (RCV) being greater than in the control period.

## Discussion

The primary goal of the study was to investigate the recovery process following an extended period of sleep restriction. We have observed differences in behavioral, motor, and neurophysiological response to both the sleep debt and the recovery. The main results are summarized in Table 1: we observe full return (reaction times in behavioral task), only partial return (actigraphy, behavioral task accuracy) or no return (ERPs, power spectrum) to the baseline values within the seven days of recovery.

Naturally, the price for allowing the participants to follow their daily routines was a number of uncontrollable factors that might have affected their lifestyle, alertness, and mood which may cause potential caveats in the interpretation of results. This further led to some subjects not being able to comply strictly with their reduced sleep schedules – such cases were monitored and revealed by actigraphy, but resulted in drop-outs. Because of that and the costs, the sample size is relatively small for an EEG study. There was also no control group, where a subject would attend 21 consecutive days of EEG measurements without any sleep restrictions. Due to the mere length and tediousness of the study one has to take into account, e.g., that in Stroop task the return to baseline accuracy might be only partial, because of the participants’ decrease in motivation. This as well invites questions about habituation effects, which are, however, addressed below. Finally, there are stable, trait-like inter-individual differences in the vulnerability to sleep deprivation^9–11^, which are not controlled in our study. On the other hand, we would like to emphasize how demanding – owing to its extended nature, subjects’ non-compliance, their increased fidgetiness having an EEG cap on, falling asleep in resting state with eyes closed, etc. – is acquisition of quality data on chronic sleep restriction.

What is novel and not previously studied is the fact that the results of this study are multi-faceted. First, the behavioral measures tend to revert to baseline values, as expected. Secondly, the actigraphy measures revert to normal only to an extent, which suggests long-lasting (even a week) disruption of both motor control and overall behaviour, including daily routines. Lastly, the EEG measures either reveal no significant change between conditions (power spectrum in task and resting state with eyes closed) or, interestingly, the change from the baseline is not reverted by the recovery (ERPs and power spectrum in resting state with open eyes).

The outcomes of power spectrum measurement require some attention, as they might seem to contradict some established results. Among the known sleep deprivation hallmarks in an increase in the theta range (4–8 Hz) power in wake EEG (open eyes)^46,47^, especially the frontal theta (closed eyes)^48^, positively correlated with subjective sleepiness during wake task EEG^49^ and with pre-existing sleep deprivation during sleep inertia period^25^, and connected, e.g., to changes in body temperature^50^. Less conveniently alpha band power was found to be either positively correlated with sleepiness (or length of wakefulness) in task EEG^49^ and resting state eyes open^46^ or negatively with eyes closed^48,51^ (the last case reportedly explained by the increased amount of microsleep ^46^). These results, however, addressed the question of acute sleep deprivation (usually, up to 40 hours of wakefulness) or a short sleep restriction (two days^52^), whose experimental designs are hardly comparable with ours. Moreover, subjects in our study had their EEG measurements at the time of day chosen according to their chronotype, and consequently close to the time of their optimal wakeful functioning – quite the opposite to what happens during prolonged wakefulness in the studies cited above.

Next, the ERP results beg the question if they might be affected by habituation to the task. The P300 amplitude – but not its latency — has been shown to habituate after repeated presentation of visual stimuli^53,54^, as the task becomes more automatic and thus attentional processes might be reduced. After several days of practice also the P300 latency can be reduced, as evidenced^55^ in a Go/No-go paradigm for No-go stimuli. At the same time P300 amplitude was shown to decrease and its latency to lengthen in response to sleep deprivation^56–58^, however, these studies dealt with acute sleep deprivation and auditory task only. On the other hand, our current result is consistent with Oginska et al.^59^. In that study, one group of subjects first went through one week sleep restriction and later, after a break, through a week of unrestricted sleep, while the second group had the order reversed. The reduction of P300 amplitude in the sleep restriction period with respect to unrestricted sleep was found irrespective of the experimental group, which rules out attributing the changes merely to habituation to repeated stimuli. Additionally, it is worth noting that the EEG resting state as well as actigraphy results are more agnostic to study design.

Present result can be explained by two not exclusive possibilities: either that after a prolonged period of sleep restriction, one week of a typical sleep routine might not be enough to fully return to the pre-experimental functioning; or that the neural activation patterns learned, and possibly optimized, during the demanding sleep conditions become consolidated. The latter – “that the brain undergoes adaptive changes in response to chronic sleep restriction, that serve to sustain a stable, albeit reduced, level of performance”^60^, and that it remains in that state – has already been suggested in previous studies^14,60^.

In the outlook, it might be worthwhile to disentangle the early and late recovery effects (the first one-two days versus a week or more) – our current sample size, however, is insufficient for robust hypothesis testing of daily measurements – and to extend the recovery period at least for the sake of actigraphy recording. We are also still aiming to look at the correlations between performance and various (reported) neurophysiological quantities, especially in the light of recently reported decline of long-range temporal correlations in the human brain during sustained wakefulness^61,62^ and the correlation of neural and behavioral scaling laws^63^. Another topic not explored here, is the topographic distribution of the changes in neural activity, which are known to differ for different tasks after total sleep deprivation^52,64^. To assess and factor out the inter- and intra-individual variability – which are known to be considerable, e.g., for alpha rhythms^65,66^ – would also involve a considerably larger, possibly a multi-center study.

## Acknowledgments

JKO was supported by the Grant 2015/17/D/ST2/03492 of the National Science Centre (Poland). TM acknowledges Grant 2011/01/B/HS6/00446 of the National Science Centre (Poland). KO acknowledges grant of Polish Ministry of Science and Higher Education 7150/E-338/M/2015 and 7150/E-338/M/2016.

## Author contributions

AD, DRC, EGN, HO, JS, MAN, MF, TM conceived of the study and participated in its design. AD, HO, JKO, JS participated in its coordination, and collected and curated the data. DRC, EGN, JKO, JS participated in designing its methodology. AMB, JKO, KO, JS participated in software development and statistical analysis. All authors participated in interpretation of the results; AMB, JKO, JS, KO drafted the manuscript. All authors read, revised and approved the final manuscript.

## Additional information

The authors declare no competing interests.

## Supplementary figures

